# Functional dissection of Wag31 domains for septal recruitment and polar distribution during the cell cycle

**DOI:** 10.1101/2025.08.21.671543

**Authors:** Julienne Petit, Daniela Megrian, Mariano Martinez, Adrià Sogues, Célia de Sousa-d’Auria, Mathilde Ben Assaya, Catherine Thouvenot, Emilie Lesur, Yann Bourdreux, Nicolas Bayan, Pedro M Alzari, Anne Marie Wehenkel

## Abstract

Bacterial cell morphogenesis is controlled by the synthesis and organization of peptidoglycan and driven by multi-protein complexes such as the divisome and elongasome. Here we investigate the role of the *Corynebacterium glutamicum* DivIVA homologue, Wag31, the elongasome scaffold essential for polar growth in *Corynebacteriales*. Conditional depletion of Wag31 results in viable but coccoid-shaped cells, showing that Wag31 is essential for rod shape maintenance. Our structural phylogenetic analyses of DivIVA homologues revealed that in *Actinobacteria*, unlike *Firmicutes*, an intrinsically disordered region spatially separates the N-terminal lipid-binding domain (LBD) from the C-terminal coiled-coil domain (CCD). We show that the LBD is necessary and sufficient for septum localization, independent of its membrane-binding properties, while the CCD domain mediates self-interaction and polar accumulation. Our findings suggest that Wag31 is recruited specifically to the septum through protein-protein interactions, priming the future pole and allowing for a timely divisome-elongasome transition at cytokinesis. Once the pole is formed the self-aggregative properties of the C-terminal CCD dominate and form a stable structure that likely organizes the pole for cell wall biosynthesis.

## Introduction

Bacterial cell shapes are driven by the synthesis of peptidoglycan (PG), a three-dimensional polymer formed by long glycan strands cross-linked via short peptide chains ^1^. Cell wall biogenesis needs to be finely regulated during the cell cycle to allow for the generation of two identical and viable daughter cells, and thus, bacterial morphogenesis relies on the tight coordination between cell division and elongation. These two processes are precisely orchestrated by large multi-protein complexes, the divisome and the elongasome respectively, that drive the assembly of the bacterial cell wall and guide PG insertion ^2^. The organization and assembly of these machineries generally rely on essential cytoskeletal proteins. Divisome assembly requires FtsZ, the nearly universally conserved tubulin homologue. However, the elongasome relies on different scaffolds in different bacteria ^3^. This diversity reflects the singular modes of bacterial cell elongation, which can proceed from the lateral side (dispersed growth), the cell centre (septo-peripheral growth) or the cell pole (polar growth) ^4^. The dispersed lateral growth, driven by the actin-like MreB protein to organise the Rod complex, has been well characterized in model bacteria like *Escherichia coli* ^5^ or *Bacillus subtilis* ^6,7^. MreB has been extensively studied and detailed molecular mechanisms underlying its function have been described ^8^. By contrast, less is known about the molecular determinants required for cell elongation in bacteria without lateral growth, which lack MreB and most of the Rod complex orthologs, except for the SEDS couple: RodA and its cognate b-PBP ^9^. This is notably the case for polar-growing rod-shaped *Corynebacteriales,* an important suborder of Actinobacteria that contains major human pathogens such as *Mycobacterium tuberculosis* or *Corynebacterium diphtheriae*. In these bacteria, the coiled-coil scaffolding protein Wag31 (DivIVA) is crucial for polar growth^10,11^.

Wag31/DivIVA is broadly conserved in Gram-positive bacteria ^12^ and in both *Firmicutes* and *Actinobacteria*, it localizes to the poles and the division site where it is thought to sense the negative curvature ^13,14^. In most *Firmicutes*, DivIVA is a non-essential protein responsible for polar localization of the MinCD complex thereby preventing the formation of the division site at the poles ^15,16^. In polar growing *Actinobacteria*, Wag31/DivIVA is an essential protein required for elongation and is thought to be the scaffold of the PG synthesis machinery ^11,17,18^. For instance, in *Streptomyces coelicolor,* DivIVA forms the polarisome that localizes to the tips of the hyphae and future sites of branching where it promotes PG synthesis ^19^. Branching appears over the lateral walls where new foci of DivIVA accumulate. Its overexpression causes hyperbranching and the tips become swollen and rounded^11^. In *Mycobacteria*, Wag31 (also known as Antigen 84) localizes asymmetrically to the cell poles as well as to the septum and this localization correlates with the incorporation of nascent PG ^18^. Indeed, PG labelling in a conditional Wag31 depletion strain showed that PG can still assemble in the absence of Wag31 but is delocalized and dispersed around the cells ^20^. Wag31 is also thought to direct the assembly of the outer layers of the complex mycobacterial cell envelope (composed of three distinct macromolecules - peptidoglycan, arabinogalactan and mycolic acids) at the pole by organizing an inner membrane domain (IMD) through direct or indirect interaction with enzymes involved in the early steps of cell wall precursor synthesis ^21,22^. In *C. glutamicum*, Wag31 overexpression leads to polar bulging and its depletion to cellular rounding ^10^. Wag31 colocalized with the SEDS GT RodA of the elongasome in cellular interaction assays ^23,24^. When expressed in *E. coli*, Wag31 and ParB also colocalize, suggesting a direct interaction with the chromosome segregation machinery ^25^.

DivIVA is a coiled-coil protein defined by its 60 amino acid N-terminal domain (DivIVA-domain) also referred to as the lipid-binding domain (LBD)^26^ as it allows to anchor DivIVA to the membrane. The LBD is followed by a C-terminal coiled-coil domain (CCD), usually flanked by intrinsically disordered regions (IDRs) of variable length ^14^. The C-terminal domain was shown to oligomerize in *B. subtilis* ^26^ and together with its low-resolution X-ray structure, suggested that the self-interaction could create a BAR-like domain ^14^ that might be responsible for negative curvature recognition, although exactly how this would work remains unclear. However almost twenty years after the first oligomers were observed in *B. subtilis*^27^, their precise structural arrangement and their role during the cell cycle remain elusive.

In this work we set out to understand the molecular function of the N- and C-terminal Wag31 domains in *Corynebacterium glutamicum*. We generated a conditional depletion strain for cellular studies and showed that this depletion, unlike what was previously described, resulted in round but viable cells that divide and that have a fully formed cell envelope. Phylogenetic studies show that Wag31 has acquired an intrinsically disordered domain between the N-terminal lipid binding domain (LBD) and the C-terminal coiled-coil domain (CCD), separating the membrane binding properties from the self-interacting properties of the protein. Using cellular localization and complementation studies we assigned distinct functional properties to the N- and C-terminal domains. First the LBD is necessary and sufficient for septum localization, likely mediated by protein-protein interactions. Second, the CCD is responsible for self-interaction and protein accumulation at the pole. Interfering with the LBD domain systematically leads to asymmetric distribution of Wag31 to one pole, suggesting that correct priming at the septum prior to cytokinesis is essential for cellular repartitioning of the elongasome scaffold. Together, these results uncover new roles and *Corynebacteriales*-specific mechanisms for this crucial protein, paving the way for detailed mechanistic studies.

## Results

### A conditional Wag31 depletion strain is coccoid shaped and viable

In *Corynebacteriales*, *wag31* is defined as an essential gene ^17^ found at the end of the division and cell wall (dcw) cluster. To understand the role of Wag31 in the cell, we designed a conditional depletion strain (*Cglu_P_ino_-wag31*) *in Corynebacterium glutamicum* (*Cglu*, ATCC13032) by placing a transcriptional terminator just before the *wag31* open reading frame, and by putting it under the control of the *myo*-inositol-repressible promoter (*P_ino_*) of the inositol phosphate synthase Ino1 gene ^28^ (Figure 1a). Wag31 protein levels were assessed after 6 hours by Western Blot revealing an almost complete depletion of the protein (Figure 1b). Surprisingly, depleted cells were viable after 24 hours of growth and kept growing in a second exponential phase in the presence of *myo*-inositol following an overnight depletion and re-dilution (Figure 1c). To investigate the impact of Wag31 depletion on cell morphology, we imaged the *Cglu_P_ino_-wag31* strain in the absence and presence of *myo*-inositol. As expected from previous work ^10,29^, depleted cells went from a rod-shaped to a coccoid morphology (Figure 1d). Despite this dramatic change in shape, the bacteria still formed septa and segregated DNA between daughter cells (Figure 1d), suggesting a fully functional divisome machinery. The depleted cells display a circularity close to 1 (0.98) (Figure 1e), their length decreases (from 2.24±0.53 μm to 1.88±0.43 μm and 1.62±0.32 μm after 6- and 24-hours depletion respectively) and width increases from 1.03±0.10 μm to 1.48±0.25 μm, which results in a mostly unchanged cell area (Figure 1e). To verify that the observed phenotype was solely due to the depletion of Wag31, we expressed Wag31 ectopically from a plasmid under the control of *P_gntK_*, a tight promoter that is repressed by sucrose (4%) and induced by gluconate (1%) ^30^. Unless otherwise stated, in the following experiments we used minimal medium (CGXII, see Materials and Methods) supplemented with 4% sucrose, where Wag31 expression is minimal (Supp. Figure S1), or 1% gluconate/4% sucrose, where Wag31 is overexpressed, and expression levels are close to native (Supp. Figure S1). The Wag31 depletion phenotype was rescued when the strain was grown for 6h in the presence of gluconate (Figure 1d,e). Circularity was 0.84±0.10, a value similar to the WT (0.82±0.08) (Figure 1e), and length and width also went back to WT-like values (Figure 1e).

**Figure 1:**
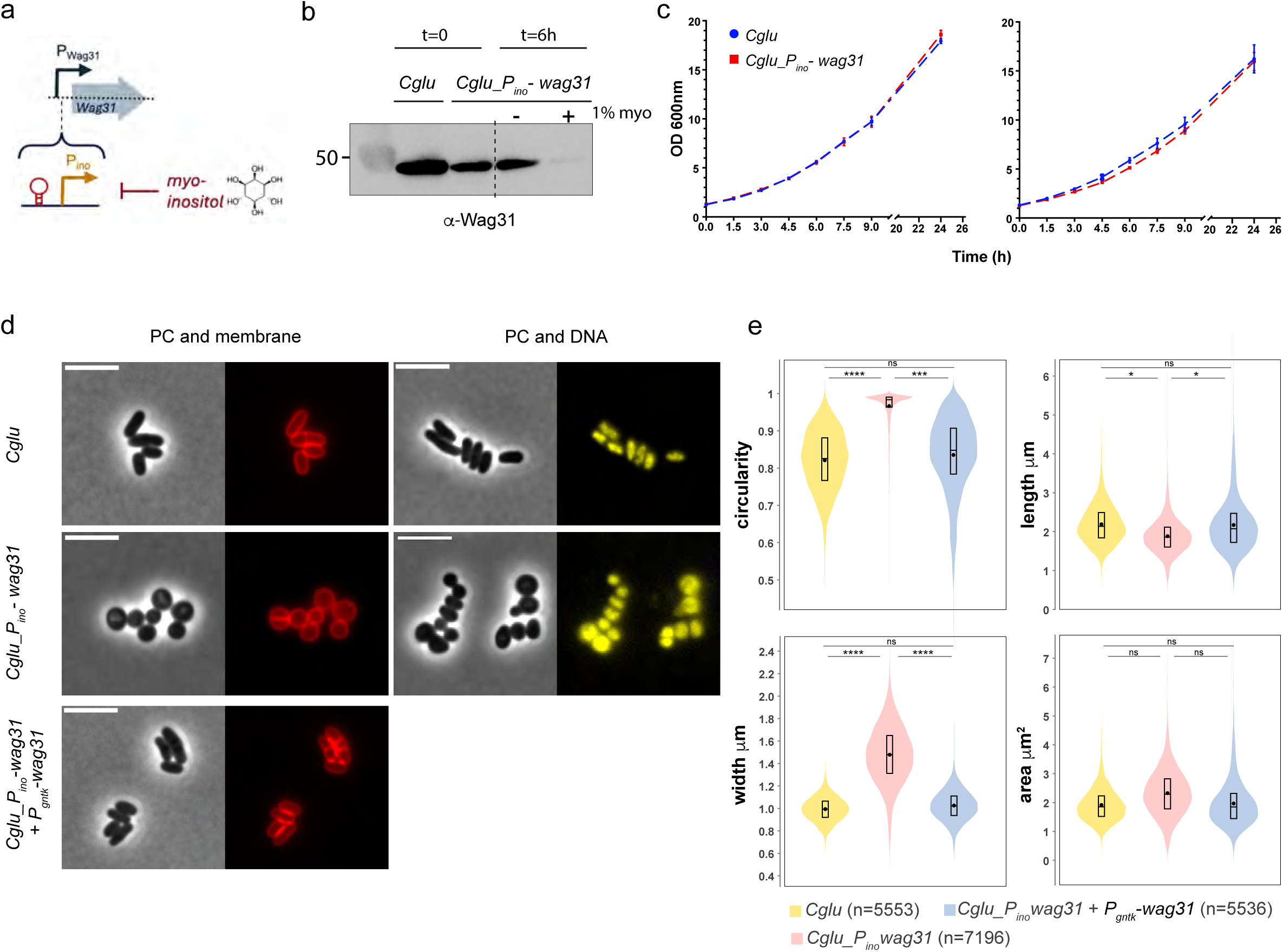
Phenotypic characterization of *Cglu_P_ino_-wag31*. **(a)** Schematic description of depletion strategy **(b)** Western blot of whole cell extracts (60 μg) from *Cglu* and *Cglu_P_ino_-wag31* after overnight growth in minimal medium (t=0) and grown for 6 hours in the absence or presence of 1% *myo*-inositol (t=6h). Wag31 levels were revealed using anti-Wag31 antibodies. **(c)** Growth curves of *Cglu* (blue) and *Cglu_P_ino_-wag31* (red) grown in minimal medium with 1% *myo*-inositol for 24 hours (left panel), then diluted a second time in fresh medium with 1% *myo*-inositol and grown for another 24 hours (right panel) **(d)** Representative images in phase contrast (PC) and Nile Red (membrane) on the left and PC and Hoechst (DNA) on the right for *Cglu*, *Cglu_P_ino_-wag31* and *Cglu_P_ino_-wag31* complemented with *P_gntK_-wag31* (+ gluconate) strains grown depleted of Wag31 for 6 hours in minimal medium **(e)** Violin plots showing the distribution of cell circularity, cell length, cell width and cell area for *Cglu*, *Cglu_P_ino_-wag31* and *Cglu_P_ino_-wag31*+ *P_gntK_-wag31* (+ gluconate) depleted for 6 hours. Cohen’s d for circularity: (^ns^d=0.22), (***d=1.79), (****d=2.46); for length: (^ns^d=0.12), (left,*d=0.76), (right,*d=0.54); for width: (^ns^d=0), (left,****d=2.26), (right,****d=2.18); for area: (top,^ns^d=0.07), (left,^ns^d=0.44), (right,^ns0.07^d=0.46).

### A full cell envelope is assembled in the absence of Wag31

To understand a possible direct role for Wag31 in regulating elongasome function and cell envelope assembly, we first assessed cell wall composition in the depleted *Cglu_P_ino_-wag31* and the *Cglu* strains. Wag31 depleted cells incorporate PG at the septum in line with a functional divisome (Figure 2a). To evaluate the integrity of the mycomembrane (MM), we stained the bacterial cells with 2-FIC5Tre^31^, a trehalose-based probe that is converted by mycoloyltransferases into a fluorescent version of trehalose monomycolate (TMM), a key glycolipid of the outer membrane. The depleted *Cglu_P_ino_-wag31* cell surface was fully stained by the probe (Figure 2b). To rule out the possible incorporation of mycolic acids before complete Wag31 depletion, we grew *Cglu_P_ino_-wag31* cells overnight in *myo*-inositol, then initiated a second exponential phase with pre-depleted cells in fresh medium to eliminate residual mycolic acids. Even under these conditions, *Cglu_P_ino_-wag31* retained an MM similar to *Cglu* (Figure 2b). Consistent with prior studies, glycolipids are incorporated after septum formation (Figure 2b, white arrows, top panel), and the uniform cell surface staining is probably due to the high fluidity of the MM of *Cglu* ^32^. To our surprise however 2-FIC5Tre staining was observed at the nascent septum (Figure 2b, white arrows, bottom panel), which differs from the previously reported WT situation where the glycolipids infiltrate the septum just before the mechanical V-snap^32^. The Wag31 depleted strains exhibited a surface similar to WT cells in scanning electron micrographs (SEM, Figure 2c). Interestingly, the rounded cells displayed deep cleavage furrows, likely marking active division sites, which differs from WT cells where characteristic holes (Figure 2c, arrows) allow for glycolipid infiltration prior to V-snapping and daughter cell separation^32^. These observations suggest that coccoid shaped cells devoid of Wag31 divide by invagination rather than mechanical V-snapping post-cytokinesis.

**Figure 2:**
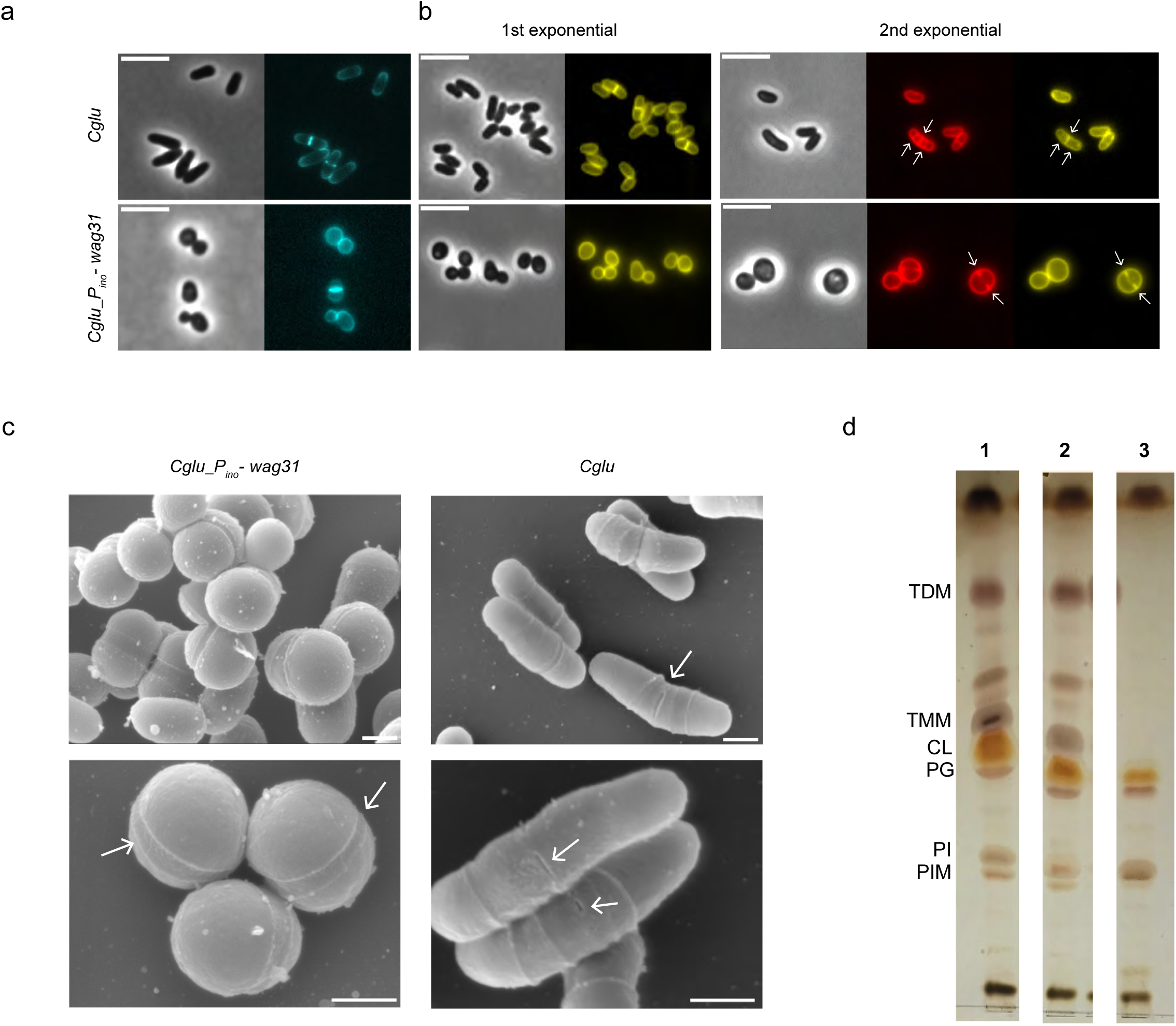
Characterization of the cell envelope of the depleted *Cglu_P_ino_-wag31* strain. **(a)** Representative images in phase contrast (PC) and HADA (PG, blue) for *Cglu* (top left) and *Cglu_P_ino_-wag31* (bottom left) depleted for 6 hours. **(b)** Representative images in phase contrast, Nile Red (membrane, red) and 2-FIC5Tre (mycomembrane, yellow) staining, for *Cglu*, and depleted *Cglu_P_ino_-wag31* in a 1^st^ and 2^nd^ (rediluted from 1^st^) culture. White arrows indicate forming septa (red channel) and TMM incorporation (yellow channel). Scale bars 5μm. **(c)** Scanning electron micrographs showing depleted *Cglu_P_ino_-wag31 (top) and Cglu (bottom)* at magnifications of 27.000x and 50.000x. Scale bars: 500nm. Arrows indicate cleavage furrows for *Cglu_P_ino_-wag31* and pre-rupture holes for *Cglu.* (d) TLC analysis of crude lipid extracts isolated from cells grown in 1% *myo*-inositol in minimal medium to stationary phase. Comparable amounts were loaded on TLC plates for Cglu (1), *Cglu_P_ino_-wag31* (2) depleted cells or Δ*pks13* (3)*. PG*, phosphatidyl-glycerol; *CL*, cardiolipin; *PI*, phosphatidylinositol; *PIM*, phosphatidylinositol mannosides; *TMM*, trehalose monomycolates; *TDM*, trehalose dimycolates.

We probed the lipid composition of the cell envelope of *Cglu* and depleted *Cglu_P_ino_-wag31* strains (Figure 2d). As a reference for affected outer membrane composition, we used a strain lacking mycolic acids (Δ*pks13* ^33^). Lipid profiles were comparable for *Cglu* and *Cglu_P_ino_-wag31*, which both show a similar signal for trehalose mono- and di-mycolates (TMM and TDM), the most abundant glycolipids of the mycomembrane generally referred to as ‘free mycolates’ because they are not covalently linked to the AG-PG layer ^34^. In contrast, the Δ*pks13* strain, which does not synthesize mycolic acids and therefore no outer membrane, has neither TDM or TMM (Figure 2d), suggesting that enzymes producing and incorporating the cell envelope are present and functional when Wag31 is depleted.

### A structural phylogenetic analysis of Wag31/DivIVA paralogues reveals major differences between Actinobacteria and Firmicutes

Wag31/DivIVA homologues display a highly conserved N-terminal lipid binding domain (LBD) or DivIVA-domain (hereafter named Wag31_LBD_) and a C-terminal coiled-coil domain (Wag31_CCD_) (^14,29^, Figure 3). Interestingly, in *Actinobacteria*, two conserved inserts can be observed between Wag31_LBD_ and Wag31_CCD_ (residues 70-175, *Cglu* numbering) and within the Wag31_CCD_ domain (residues 285-318), making these paralogs much longer than in other phyla (Figure 3). Structural studies of the full-length Wag31 are challenging as the protein tends to aggregate and form gel-like material during purification. This gel could be partially solubilized at high pH and high salt concentrations, but unsurprisingly the protein did not yield protein crystals, and did not result in interpretable images in cryo-EM. To obtain structural information on the full-length protein we used AlphaFold (AF) to compute structural models of dimeric full-length Wag31 from *C. glutamicum* and *M. tuberculosis*, and DivIVA from *B. subtilis* (Figure 4a). Interestingly Wag31 models suggest that the LBD domain is separated from the CCD by a disordered region that corresponds to the *Actinobacteria*-specific insert (Figure 3). In contrast, the structure of *B. subtilis* DivIVA is predicted to form a continuous coiled-coil extending from the N-terminal LBD-domain into the C-terminus (Figure 4a). To understand if this is a conserved feature of Firmicutes we calculated structural models of representative DivIVA dimers for each order on the schematic phylogenetic tree shown in Figure 4b. Strikingly, the evolutionary history as well as the sequence alignment and structure predictions of the represented DivIVA proteins indicate that the coiled-coil domain from the Firmicutes DivIVA-domain forms a single, continuous coil, whereas actinobacterial proteins (except for the most ancestral one, that may represent a transition) present an insert between the LBD and the CCD spatially separating the former from the latter (Figure 4c). This acquired flexibility could allow for multiple conformational states and might explain the dynamic and functional differences between DivIVA in Firmicutes and Actinobacteria ^35^.

**Figure 3:**
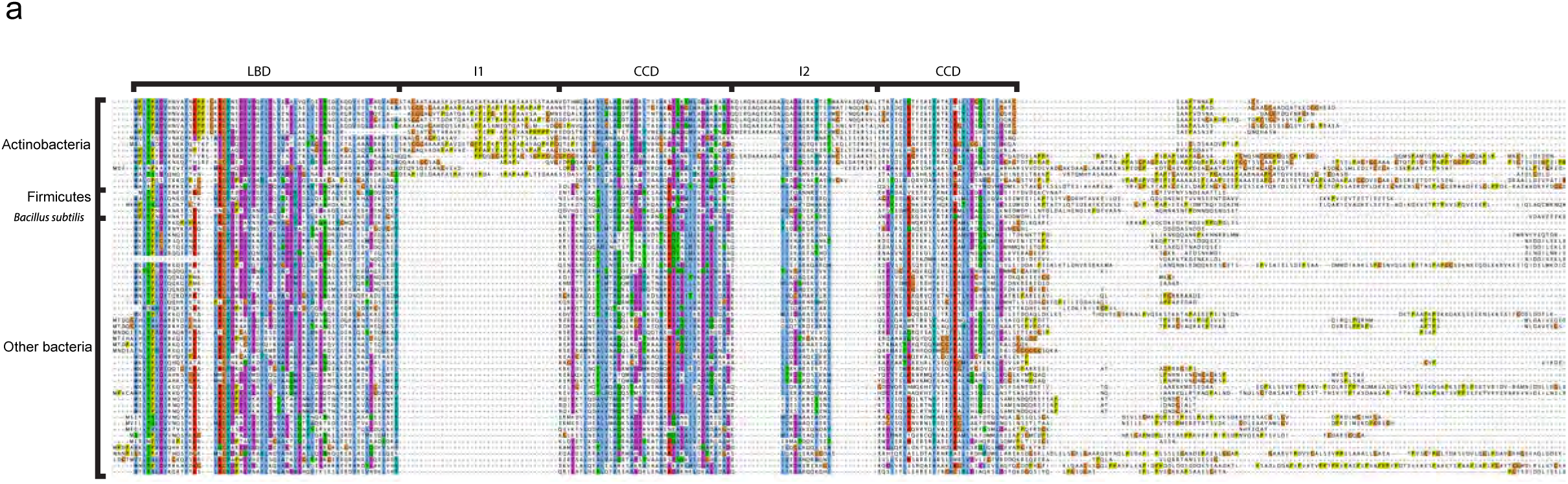
DivIVA/Wag31 family domain definition from sequence alignment. Trimmed multiple sequence alignment of DivIVA/Wag31 in all Bacteria. Conserved domains (LBD and CCD) and inserts (I1 and I2) are indicated at the top. Sequences of Firmicutes, B. subtilis and of Actinobacteria are indicated on the left.

**Figure 4:**
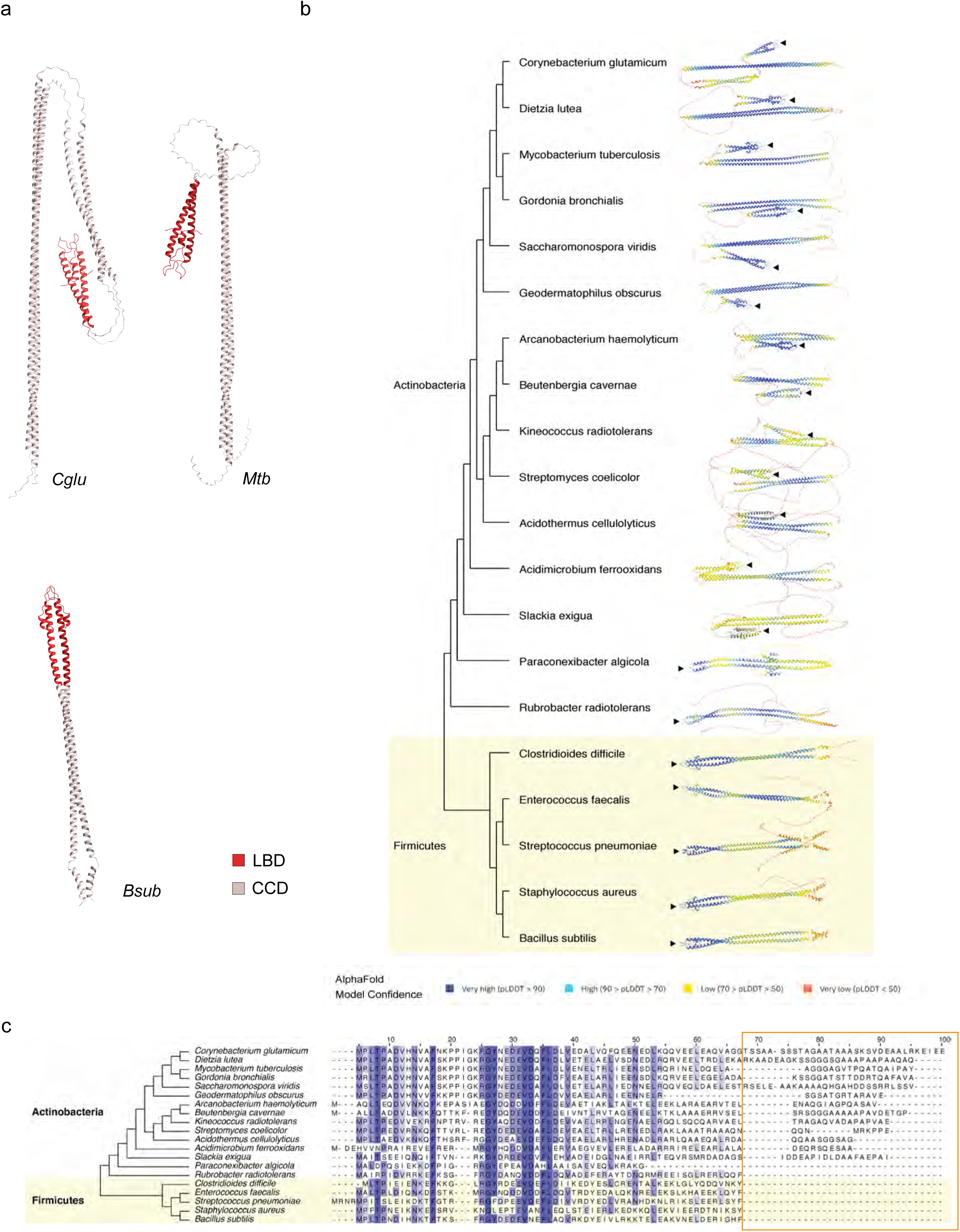
Structural and phylogenetic analysis of DivIVA/Wag31 domains. **(a)** Alphafold prediction for dimeric full-length Wag31 homologues from *Cglu, Mtb,* and *Bsub*. **(b)** Alphafold prediction for dimeric full-length Wag31 of representative species of Actinobacteria and Firmicutes colored according to the Alphafold model confidence and associated schematic phylogeny. The black arrow indicated the membrane binding tip of the LBD **(c)** Schematic phylogenetic tree and multiple sequence alignment showing the sequences of the same representative species as in (b) and highlighting the insertion after the DivIVA-domain present in *Actinobacteria* (orange frame).

### Distinct functional properties of the Wag31 N- and C-terminal domains

To understand the specific roles of Wag31_LBD_ and Wag31_CCD_, we expressed both truncation mutants in the *Cglu_P_ino_-wag31* strain. The *wag31* depletion phenotype was not rescued when the strain was complemented with Wag31_LBD_-mNeon (Figure 5a). Cells were round and displayed similar circularity (0.95) than the depleted *Cglu_P_ino_-wag31* strain. Importantly, Wag31_LBD_ could still localize to the septum, where PG incorporation occurs (Figure 5a), pointing to Wag31_LBD_ as a driving force for septal localization of the full-length protein. To investigate whether this interaction depends on the membrane binding properties of the Wag31_LBD_ we made a triple point mutant (I18D, K20S, R21S) in the membrane binding tip of the LBD domain (Wag31_LBD_mut_). This mutant construct retained its septal localization, indicating that membrane binding is not essential for Wag31 recruitment to the division site (Figure 5b) and suggesting that this recruitment is instead mediated by protein-protein interactions. We have previously shown that the membrane-associated Glp-GlpR complex mediates Wag31 recruitment to the mid-cell divisome *in vivo* ^36^ and that Wag31_LBD_ directly interacts with GlpR *in vitro*, with an apparent dissociation constant in the micromolar range^36^. The interaction between GlpR and Wag31_LBD_ could thus be a major force behind divisome recruitment and septal localization of Wag31.

**Figure 5:**
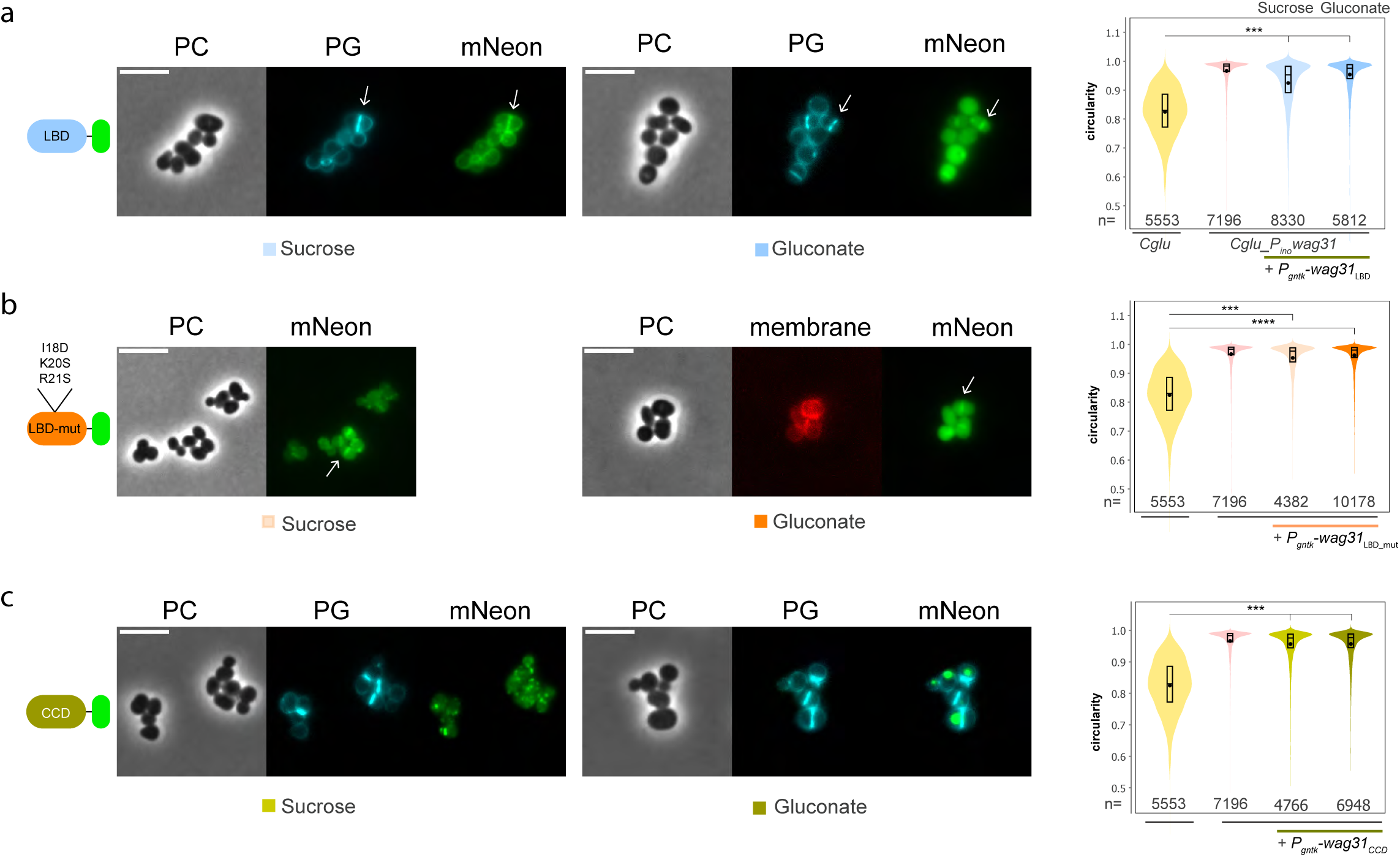
Domain characterization of Wag31_LBD_ and Wag31_CCD_. **(a)** Representative images in phase contrast (PC), HADA (PG) and green fluorescent signal (mNeon) in depleted *Cglu_P_ino_-wag31 + P_gntK_-Wag31_LBD_-mNeon*. Low and high expression levels of Wag31_LBD_-mNeon in sucrose and gluconate, respectively. White arrows indicate septal PG and septal localization of Wag31_LBD_-mNeon. Violin plots showing the distribution of cell circularity in *Cglu* (WT), depleted *Cglu_P_ino_-wag31* (Pino 6h) and depleted *Cglu_P_ino_-wag31* expressing Wag31_LBD_-mNeon in sucrose without or with 1% gluconate (labelled sucrose or gluconate, respectively). Cohen’s d: (***, 1.25, 1.96). **(b)** Representative images in phase contrast (PC), HADA (PG) and green, fluorescent signal (mNeon) in depleted *Cglu_P_ino_-wag31 + P_gntK_-Wag31_LBD-mut_-mNeon*. Low and high expression levels of Wag31_LBD-mut_-mNeon in sucrose and gluconate, respectively. White arrows indicate septal localization of Wag31_LBD-mut_-mNeon. Violin plots showing the distribution of cell circularity in *Cglu*, depleted *Cglu_P_ino_-wag31* and depleted *Cglu_P_ino_-wag31* expressing Wag31_LBD-mut_-mNeon in sucrose without or with 1% gluconate. Cohen’s d: (***, 1.81) (****, 2.25). **(c)** Representative images in phase contrast (PC), HADA (PG) and green, fluorescent signal (mNeon) in depleted *Cglu_P_ino_-wag31 + P_gntK_-Wag31_LBD-mut_-mNeon*. Low and high expression levels of Wag31_CCD_-mNeon in sucrose and gluconate, respectively. White arrows indicate septal localization of Wag31_CCD_-mNeon. Violin plots showing the distribution of cell circularity in *Cglu*, depleted *Cglu_P_ino_-wag31* and depleted *Cglu_P_ino_-wag31* expressing Wag31_CCD_-mNeon in sucrose without or with 1% gluconate. Cohen’s d: (****, 2.06, 2.15). All scale bars 5μm.

Ectopic expression of Wag31_CCD_-mNeon in the depleted *Cglu_P_ino_-wag31* strain also failed to complement (Figure 5c), but led, in a concentration dependent manner, to a single aggregative spot randomly localized in the cell (Figure 5c). The Wag31_CCD_-mNeon construct neither localized to the septum nor did it show any membrane localization but aggregated as a single spot in the cell. This suggests that the Wag31_CCD_ domain is primarily responsible for self-interaction and network formation at the poles of the WT cells, whereas Wag31_LBD_ is mainly responsible for specific divisome interactions and septum localization. Although the structural basis of coiled-coil-mediated protein oligomerization in *Corynebacteriales* remains to be determined, one possibility could be that the parallel coiled-coil dimers might associate into tetramers in which two dimers are joined in the middle by a short anti-parallel four-helix bundle, as suggested by high-confidence AlphaFold predictions (Suppl Fig. S2) and resembling similar arrangements previously seen in the low-resolution structure of *B. subtilis* DivIVA ^14^. However, such interactions cannot account for the higher order protein oligomers that likely occur at the poles. Indeed, higher order oligomers were described to form in mycobacterial cell extracts and that overexpression of Wag31 leads to overaccumulation at one pole ^37^. The additional oligomeric interactions may be driven by other regions of the C-terminal CCD or the IDRs and the role of additional protein factors in Wag31 polar aggregation cannot be excluded.

### An altered Wag31_LBD_ domain leads to Wag31 polar asymmetry

To understand the role of the LBD and CCD in cellular distribution of Wag31, we expressed different full-length constructs in the *Cglu_P_ino_-wag31* or *Cglu* strains. Heterologous expression of Wag31-mNeon, mNeon-Wag31 or a mutant in the membrane binding tip (Wag31_mut_-mNeon) in the *Cglu_P_ino_-wag31* strain restored rod-shape (Figure 6a). However, only the C-terminally tagged Wag31-mNeon phenocopied the slightly asymmetric Wag31 localization to the poles and to the septum that was observed when Wag31-mNeon was expressed in the WT (Figure 6b). In contrast, both the N-terminal mNeon fusion and the LBD point mutations in the C-terminal mNeon fusion led to a strong asymmetric polar distribution in *Cglu_P_ino_-wag31* (Figure 6a), suggesting that steric hindrance by mNeon at the Wag31 N-terminus or altered membrane-binding properties interfere with correct cellular distribution of Wag31 in the absence of the endogenous protein. In the WT *Cglu* strain where endogenous Wag31 is present, the heterologous expression of mNeon-Wag31 does not significantly alter polar and septal localization (Figure 6b). However, the expression of Wag31_CCD_-mNeon (lacking the LBD) or Wag31_mut_-mNeon (with an impaired LBD for lipid binding) led to a striking unipolar localization and very little signal at the septum (Figure 6b), suggesting that a functional membrane-binding domain is crucial for correct cellular distribution. As the membrane-binding function of the Wag31 LBD is not required for septal recruitment, the observed asymmetry likely results from perturbed dynamics and cellular equilibria, where the self-interacting properties of the CCD dominate over septal recruitment when membrane binding is impaired.

**Figure 6:**
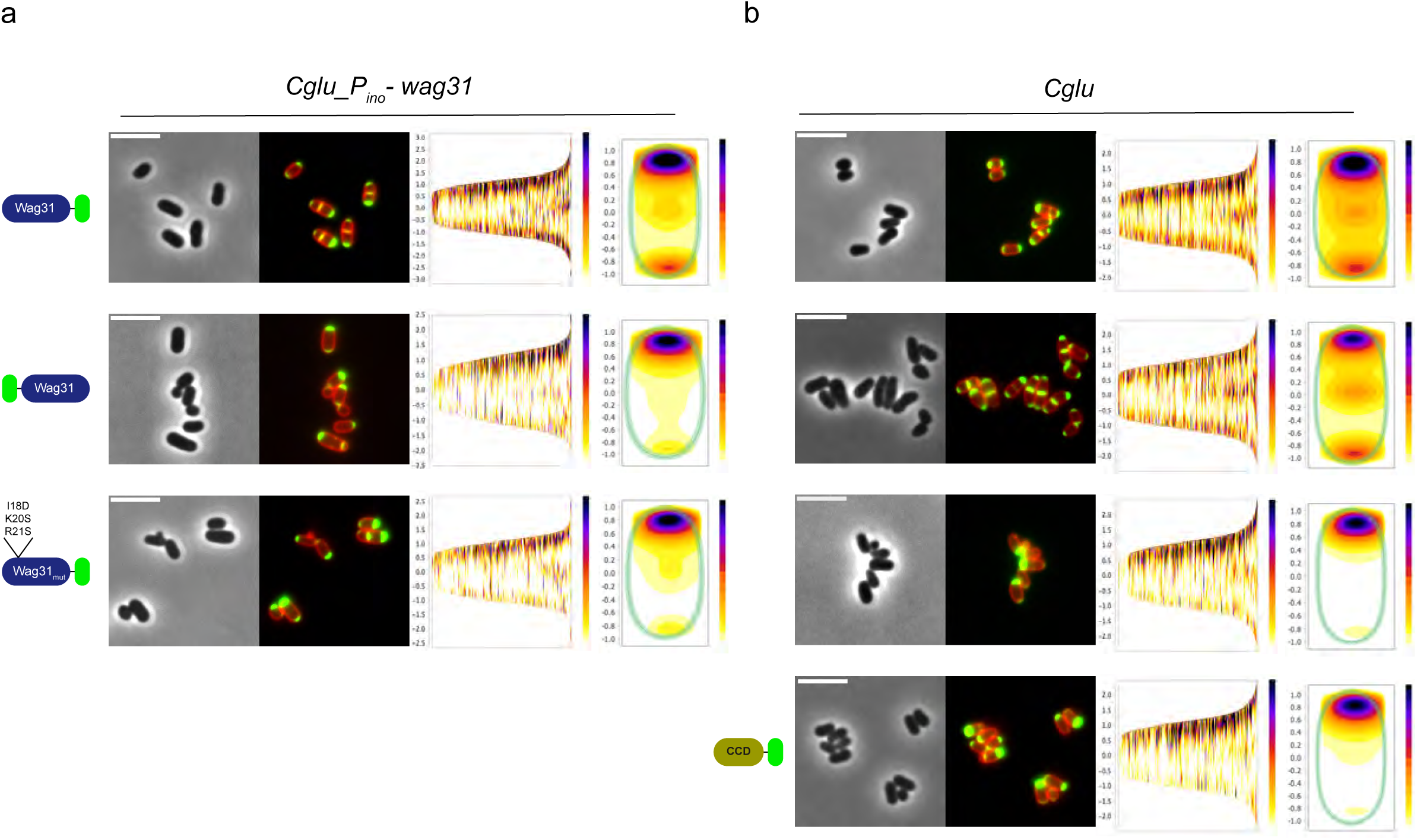
Asymmetric Wag31 localization when the LBD domain is compromised. **(a)** Representative images in phase contrast and superposed Nile Red and mNeon fluorescent signal in the depleted *Cglu_P_ino_-wag31* strain overexpressing the different constructs represented on the left. Cell profile alignments (cells sorted by length) and heatmaps representing the localization pattern of each construct. **(b)** Representative images in phase contrast and superposed Nile Red and mNeon fluorescent signal in the *Cglu* strain overexpressing the different constructs represented on the left. Cell profile alignments (cells sorted by length) and heatmaps representing the localization pattern of each construct. All cells were grown in minimal medium with 1% gluconate, +/− 1% *myo*-inositol. All scale bars 5μm.

In all the constructs that induce a unipolar localization of Wag31, the septum is often misplaced, from a mid-cell localization towards the Wag31-free pole (Figure 7a, white arrows). Quantification of septum position in the *Cglu* strain expressing either Wag31_CC_-mNeon or Wag31_mut_-mNeon revealed two populations of cells: one with a septum close to mid-cell (relative position close to 0.5, where the pole with Wag31 accumulation is taken as position 0), and one with a septum close to the pole without Wag31 accumulation (relative position around 0.75-0.8) (Figure 7b). This contrasts with the single population at a relative position close to 0.5 observed under Wag31-mNeon overexpression (Figure 7b). Unipolar localization also showed defects in DNA segregation leading to round mini-cells devoid of DNA (Figure 7a, white arrows), likely generated after division of misplaced septa. Thus, a unipolar localization of Wag31 because of a dysfunctional LBD could interfere with correct chromosome segregation.

**Figure 7:**
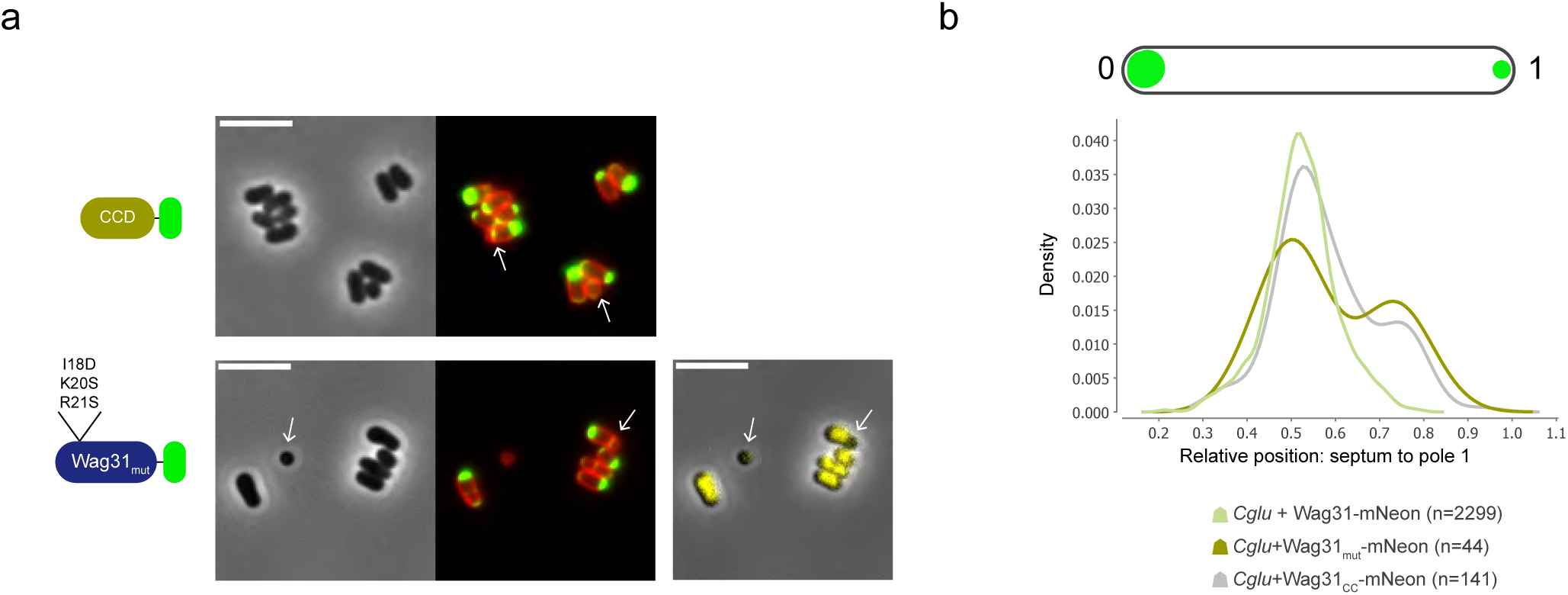
Unipolar localization leads to septum misplacement and creates a population of mini-cells. **a)** Representative images in phase contrast, Nile Red and mNeon fluorescent signal (red/green overlay), and DNA stain (Hoechst) shown in yellow in *Cglu* overexpressing the different constructs represented on the left (in minimal medium with 1% gluconate). White arrows indicate mini-cells and misplaced septa. **b)** Quantification of septum position in the *Cglu* strain expressing either Wag31_CCD_-mNeon or Wag31_mut_-mNeon highlighting the population of cells with a septum far from the brightest pole (which is position 0). All scale bars 5μm.

## Discussion

In this study we addressed domain functionality of Wag31 (DivIVA), a cytoskeletal protein that is largely documented but mechanistically poorly understood. Our structural phylogenetic analysis of DivIVA homologues revealed a clear cutoff in sequence and structure between Firmicutes and Actinobacteria, which correlates with the different functions that have been associated with DivIVA-like proteins ^12^ and could explain why DivIVA from the Actinobacteria *S. coelicolor* or *M. tuberculosis* rescued the morphological defect of a *C. glutamicum* strain depleted of Wag31 and restore polar PG synthesis, unlike DivIVA from the Firmicutes *B. subtilis* or *S. pneumoniae* ^10^. The molecular break in the form of an intrinsically disordered region which appeared very early in the actinobacterial tree, separates the membrane/septum binding function from the self-interacting coiled-coil function. This separation renders the protein more amenable to conformational regulation, by for example regulating the accessibility of the coiled-coil domain, which could in turn guide the dynamic properties and the septal/polar distribution of the protein, as described below. This evolutionary insight has led us to probe in more detail the molecular function of each domain and its implication for cell cycle progression.

### Assignment of specific molecular functions to different regions of Wag31

We have shown that the LBD is a major determinant for septal recruitment likely through protein-protein interactions with GlpR ^36^, RodA ^24^ and probably other - yet to be identified - partners. Indeed, in Firmicutes the DivIVA domain of GpsB paralogs have been shown to directly interact with penicillin-binding proteins (PBPs) in different species ^38,39^ and to form a hub for protein-protein interactions at the septum. The amino acids responsible for membrane binding, although not essential for septal recruitment, are required for correct cellular distribution of Wag31, in line with previous work where a single K20 mutant expressed in *Cglu* ^24^ or *Mycobacterium smegmatis* ^40^ failed to correctly distribute to the poles.

On the other hand, the C-terminal CCD of Wag31 primarily mediates protein self-assembly at the cell poles and may be considered the cytoskeletal domain of the protein. Indeed, Wag31 has a high tendency to self-interact and aggregate through its coiled-coils: the full-length protein forms a gel-like material *in vitro*, fluorescently labelled Wag31 forms very bright foci at the poles, and the CCD alone forms single, randomly localized, aggregative spots in depleted *Cglu_P_ino_-wag31* cells. Interestingly, when endogenous Wag31 is present, the CCD accumulates asymmetrically at one pole only.

### Driving forces for Wag31 cellular distribution and function

Coiled-coil mediated self-interaction likely drives Wag31 accumulation, acting as a template for further self-assembly. If Wag31 is initially recruited to the septum via the LBD-domain, this septal recruitment could act as a priming mechanism to nucleate Wag31 at the new poles of daughter cells upon cytokinesis. In Firmicutes the main driving force behind Wag31 localization has been attributed to its recognition of negatively curved membranes, although how this would work, from a molecular point of view, is not clear ^13,14,41^. In *Corynebacteriales*, Wag31 localization appears to be minimally influenced by membrane curvature, as shown by the strong asymmetrical localization of Wag31 variants such as mNeon-Wag31 or Wag31_mut_-mNeon. These constructs do not equally recognize both cell poles and exhibit reduced septal localization, even if all these regions are negatively curved. Moreover, the Wag31_LBD_ domain alone localizes to the septum independently of the membrane. We propose that Wag31 localization in *Corynebacteriales* is instead governed by two key mechanisms: (1) protein-protein interactions that initiate the dynamic recruitment of Wag31 to the septum, and (2) auto-accumulation in a gradient-dependent manner, that promotes further localization to regions where it is already enriched (Figure 8). Balance between these mechanisms is essential for proper bipolar and septal localization of Wag31. Disrupting this equilibrium, by interfering with either mechanism, results in aberrant distribution. This model would explain the observed asymmetry when the LBD is altered. Mutations in the LBD, steric hindrance, or domain deletion reduce Wag31 recruitment to the septum, shifting the localization equilibrium towards one cell pole.

**Figure 8:**
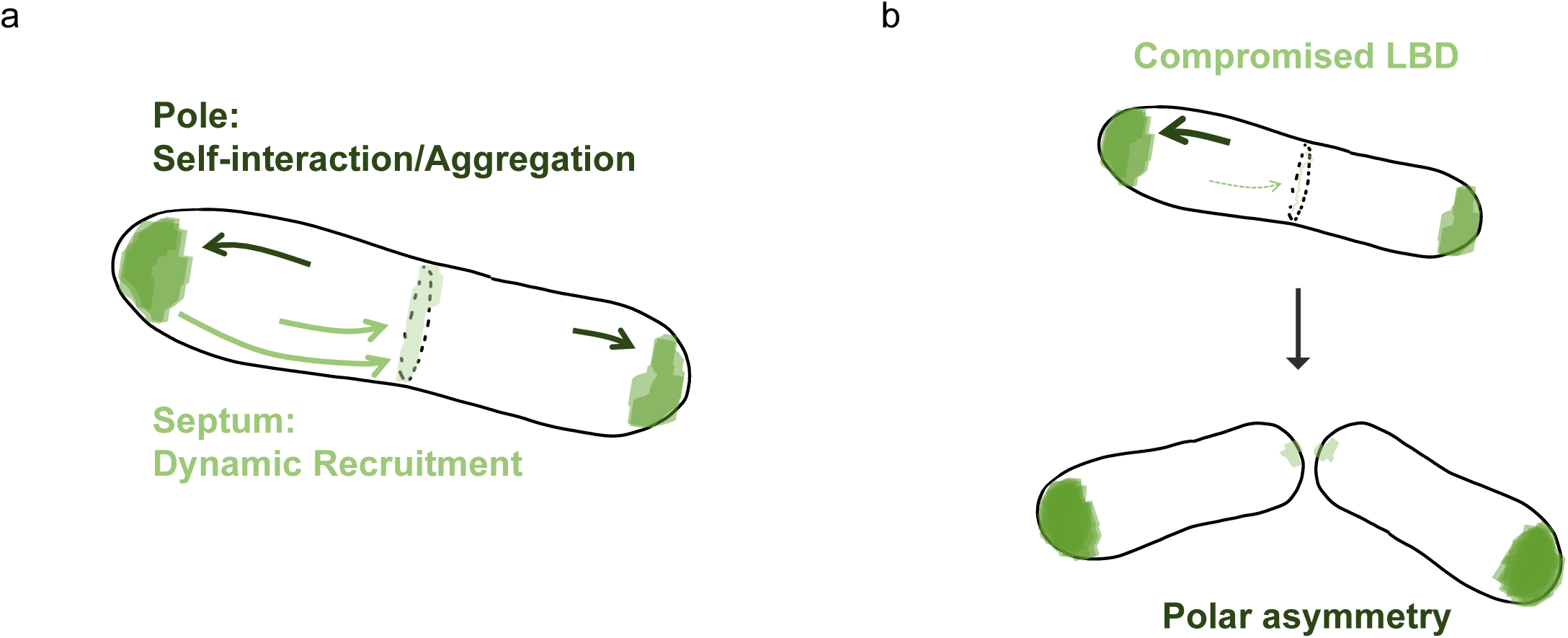
Working model for Wag31 localization in wild-type (WT) and LBD-perturbed cells. **(a)** In the WT, Wag31 is dynamically recruited to the septum, via direct protein-protein interactions mediated by its LBD domain. This serves as a nucleation for the future pole. Upon cytokinesis, Wag31 accumulates at the poles, most likely driven by coiled-coil mediated self-interaction. **(b)** Disruption of the LBD—for example, by expressing the CCD alone, fusing mNeon to the N-terminus, or introducing mutations in the lipid-binding tip—leads to aberrant cellular distribution of Wag31, resulting in strong asymmetry. This suggests that proper LBD function is required for redistribution and spatial regulation of Wag31.

The above model emphasizes the importance of understanding Wag31 dynamics to elucidate its function. Given the critical role of the LBD in septal localization, its accessibility likely influences the balanced distribution of Wag31 between the septum and the poles, and consequently between the two poles post-division. The different protein mobility observed between *B. subtilis* DivIVA and *Cglu* Wag31, with the latter being more confined to the poles ^35^, may stem from differential conformational flexibility and LBD accessibility between *Firmicutes* and *Actinobacteria*, which could alter the accessibility of the coiled-coil domain. In agreement with previous observations ^35^, our results indicate that there could be distinct Wag31 states within the cell: a dynamic state capable of migrating to the septum, and a more inert state consisting of higher-order oligomers or aggregates at the poles (Figure 8).

### Viability, morphology and cell envelope biogenesis

Under the conditions used in this study, cells depleted for Wag31 were viable in line with a recent report describing a transposon insertion in the *wag31* gene in *Cglu* ^42^. This was surprising as several past reports indicated that Wag31 is an essential gene in *Cglu* ^17^, as well as in *Streptomyces coelicolor* ^11^, *Mycobacterium tuberculosis* ^43^, and *Mycobacterium smegmatis* ^18^. Although Wag31-depleted cells remain viable, capable of dividing and assembling a complete cell envelope, they exhibit significant morphological changes, transitioning from a rod to a coccoid shape. This indicates that Wag31 plays a crucial role in maintaining cell shape and can thus be classified as a morphogenic protein. The rod-to-coccus transition upon Wag31 depletion is likely a result of disrupted polar peptidoglycan (PG) synthesis and regulation. Interestingly, *wag31* depleted coccoid cells can form a septum where they segregate their DNA and synthesize PG and the outer cell envelope components. However, for proper cell division to occur, it seems better to have no Wag31 than to have one with a corrupt LBD, as complementation of *wag31* depletion with constructs containing a dysfunctional LBD led to a strong asymmetry, with unipolar localization, misplaced septa and anucleate minicells. Wag31 was previously shown to interact with ParB in *Cglu*, which in turn forms the ParB-*oriC* nucleoprotein complex that tethers the chromosome to the cell pole for correct segregation ^25^. Therefore, a unipolar localization of Wag31 or a dysfunctional LBD could interfere with correct chromosome segregation. It is plausible that the polar Wag31 foci attract and confine ParB-*oriC* - and thus the chromosomes - to one pole only, as the opposite pole has much less Wag31 and would fail to retain the sister *oriC*. As septum placement in the cell was also found to depend on chromosome segregation, through potential nucleoid associated factors or just steric considerations ^44^, division site selection would occur where there is no DNA, moving away from the pole where Wag31 accumulates (Figure 7).

Together our work suggests that *actinobacterial* Wag31 may be regulated differently from *Firmicutes* DivIVA throughout the cell cycle, in line with its different biological functions. Driving forces other than affinity for negatively curved membranes exist to promote a balanced polar and septal localization, including species-specific interactions with the LBD. Specific regulation of Wag31 oligomerization via protein-protein interactions with other divisome components and/or post-translational modifications might prevent the premature formation of polar-like aggregates at the septum through conformational or electrostatic control. The dynamic interplay of these different elements during the cell cycle, together with putative interactions with additional regulatory factors yet to be discovered, will ultimately define the molecular properties and cellular roles of actinobacterial Wag31 in cell morphogenesis and polar growth.

## Materials and Methods

All bacterial strains, plasmids and primers used in this study are listed in Supplementary Table S1.

### Bacterial strains and growth conditions

*Escherichia coli* DH5α or CopyCutter EPI400 (Lucigen) strains were used for cloning and grown in Luria-Bertani (LB) broth or agar plates at 37 °C supplemented with 50 μg/ml kanamycin when required. *C.* glutamicum ATCC 13032 (*Cglu*) was used as a wild-type (WT) strain. *Cglu* strains were grown in brain heart infusion (BHI) or CGXII minimal medium ^45^ plus 4% sucrose at 30°C and 100-120 rpm shaking, supplemented with 25 μg/ml kanamycin when required. When specified, CGXII medium was supplemented with 1% gluconate and/or 1% *myo*-inositol.

### Cloning for protein expression in *Cglu*

For ectopic recombinant expression of the different constructs in *Cglu*, we used the pUMS_3 shuttle expression vector ^46^, in which the gene of interest is placed under the control of *P_gntK_*, a tight promoter that is repressed by sucrose and induced by gluconate ^30^. Wild-type or mutant versions of *wag31* were assembled in this plasmid by either Gibson assembly, domain deletions by PCR or site-directed mutagenesis using the primers listed in the Table S1.

For cellular localization studies, codon optimized mNeonGreen was ordered from Genscript and cloned alone (pUMS_17) or fused in frame to the N-terminus (pUMS_21) or C-terminus (pUMS_25) of Wag31 wild-type or mutants constructs, including a LEGSGQGPGSGQGSG flexible linker between the two fused proteins. The plasmids pUMS_175 and pUMS_176 were obtained by deletion of the regions of interest using the plasmid pUMS_25 as a template. The plasmids pUMS_254 and pUMS_255 were generated by site-directed mutagenesis using plasmids pUMS_25 and pUMS_175, respectively, as templates.

### *Cglu_P_ino_-wag31* conditional depletion strain construction

The conditional depletion strain *Cglu_P_ino_-wag31* was generated using a two-step allelic exchange strategy with the suicide plasmid pK19mobsacB ^47^. In brief, to achieve regulated expression of the *wag31* gene, a transcriptional terminator was inserted upstream of *wag31*, followed by the native promoter of the *ino1* gene (*cg3323*), which is repressible in the presence of *myo*-inositol ^28^. The transcriptional terminator and *ino1* promoter fragment were PCR-amplified from the plasmid pK19-P3323-lcpA7 ^48^. Approximately 500 bp regions upstream and downstream of *wag31* were amplified from the genomic DNA of *Cglu*. All DNA fragments were assembled into the pK19mobsacB backbone using Gibson assembly (New England Biolabs).

The plasmid was sequenced and electroporated into *Cglu*. Positive colonies were grown in BHI media supplemented with 25 µg/ml kanamycin overnight at 30 °C and 120 rpm shaking. The second round of recombination was selected by growth in minimal medium CGXII plates containing 10% (w/v) sucrose. The insertion of the *ino1* promoter was confirmed by colony PCR and sequenced (Eurofins, France). All the oligonucleotides used to obtain and check this strain are listed in Table S1.

### Western Blots

To prepare cell extracts, bacterial cell pellets were resuspended in lysis buffer (50 mM Bis-Tris pH 7.4; 75 mM 6-Aminocaproic Acid; 1 mM MgSO_4_; Benzonase and protease Inhibitor) and disrupted at 4°C with 0.1 mm glass beads and using a PRECELLYS 24 homogenizer. Total extracts (60 μg) were run on an SDS-PAGE gel, transferred onto a 0,2 μm nitrocellulose membrane and incubated for 1h with blocking buffer (5% skimmed milk in TBS-Tween buffer (Tris-HCl pH8 10 mM; NaCl 150 mM; Tween20 0,05% (vol/vol)) at room temperature (RT). Blocked membranes were incubated for 1h at RT with the anti-Wag31 antibody ^36^ diluted in blocking buffer (1:500). After washing in TBS-Tween, membranes were probed with an anti-rabbit horseradish peroxidase-linked secondary antibody (GE healthcare) for 45 minutes. For chemiluminescence detection, membranes were washed with TBS-Tween and revealed with HRP substrate (Immobilon Forte, Millipore). Images were acquired using a ChemiDoc MP Imaging System (Biorad).

### Phase contrast and fluorescence microscopy

For imaging, cultures of *Cglu* were grown in BHI for around 6 hours (except for *Cglu_P_ino_-wag31* strain, which was directly inoculated in CGXII 4% sucrose medium), then pelleted at 5200 x g at room temperature and inoculated into CGXII media supplemented with 4% sucrose and kanamycin (25 μg/ml) for overnight growth. The following day, cultures were diluted to an OD_600_ of 1 in CGXII supplemented with 4% sucrose (with or without 1% gluconate, 1% *myo*-inositol) and grown for 6 hours to a required OD_600_ of about 4-6 (early exponential phase). For each sample, 100 μl of cultures were pelleted, washed with fresh medium and diluted to an OD_600_ of 3 for imaging. For PG labeling, HADA stain (Tocris) was added to the culture at a final concentration of 0.5 mM, incubated for 20 min at 30 °C and 120 rpm protected from light, and washed 2 to 3 times. For MM labeling, 2-Fic5Tre dye ^31^ was added to the culture at 33 μM final concentration, incubated for 5 hours at 30 °C and 120 rpm and washed 4 times. For DNA staining, Hoechst stain was added to the culture at a final concentration of 2 ug/ml. For membrane staining, Nile Red (Enzo Life Sciences) was added to the culture (2 μg/ml final concentration) just prior to placing them on 2% agarose pads prepared with the corresponding growth medium. Cells were visualized using a Zeiss Axio Observer Z1 microscope fitted with an Orca Flash 4 V2 sCMOS camera (Hamamatsu) and a Pln-Apo 63X/1.4 oil Ph3 objective. Images were collected using Zen Blue 2.6 (Zeiss), segmented with a custom trained version of Omnipose ^49^ and analyzed using the software Fiji ^50^ and the plugin MicrobeJ ^51^ to generate violin plots and fluorescent intensity heat maps. Because of the important number of cells analyzed in each sample, Cohen’s *d* value was used to describe effect sizes between different strains independently of sample size:

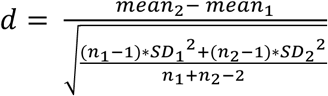

Values were interpreted according to the intervals of reference suggested by Cohen ^52^ and expanded by Sawilowsky ^53^, as follows: very small and small (ns), *d* < 0.5; medium (*), 0.5 < *d* < 0.8; large (**), 0.8 < *d* <1.2; very large (***), 1.2 < *d* < 2; huge (****), *d* > 2.

### Scanning electron microscopy (SEM)

*Cglu* and *Cglu_P_ino_-wag31* were grown as described above, harvested in early exponential phase (OD_600_ around 4). The cells were fixed in 2.5% glutaraldehyde overnight at 4°C, and postfixed for 1h in 1% osmium. Then, samples were dehydrated through a graded ethanol series followed by critical point drying with CO_2_. Dried specimens were gold/palladium sputter-coated with a gun ionic evaporator PEC 682. The samples are imaged in a JEOL IT700HR field emission scanning electron microscope.

### Sequence analyses and structure prediction

We assembled a proteomes database containing representative species of all Bacteria obtained from ^54^ (excluding Candidate phyla), five *Firmicutes* model organisms, and 15 representative species covering all actinobacterial diversity (see Data Availability). We used HMM profile searches to identify protein Wag31 in the protein database. First, we used the HMMER package (v3.3.2) ^55^ tool jackhmmer to look for homologs of *Cglu* Wag31 in all the proteomes using the GenBank ^56^ sequence BAB99543.1 as query. The hits were aligned with linsi, the accurate option of mafft (v7.475) ^57^ and default parameters. The alignments were manually curated, removing sequences that did not align globally. The hits obtained by jackhmmer might not include sequences that are very divergent from the single sequence query. For this reason, the alignment was used to create an HMM profile using the HMMER package (v3.3.2) tool hmmbuild. This specific and curated HMM profile of Wag31 was used for a second and final round of searches against the proteomes using the HMMER tool hmmsearch. The new hits were aligned with linsi, the accurate option of mafft (v7.475), and trimmed using trimal (v1.4.rev15) ^58^ to remove the columns with more than 90% of gaps to facilitate the visualization.

We predicted the structure of the dimeric form of all Wag31 homologs identified in Actinobacteria and Firmicutes using Alphafold ^59^. We used software ChimeraX ^60^ to map the Alphafold model confidence on the predicted structures.

### TLC-Cell envelope analysis

Lipids were extracted from wet pellets of cells (grown in stationary phase) with CH_3_OH/CHCl_3_ (2:1 v/v) at room temperature for 16h. The extract was centrifuged for 10 min at 6000g to recover the supernatant. After evaporation of the organic phase, the lipids were solubilized once in 1 ml CHCl_3_ and dried again. Finally, the lipid extract was resuspended in CHCl3 (generally 50 μl for 20 ml of cells in stationary phase) and then analyzed by thin layer chromatography on silica gel (TCL G-60 glass plates, 0.25 mm thick, Macherey-Nagel) developed with CHCl_3_/CH_3_OH/H_2_O (65:25:4, v/v/v). Lipid detection was performed by immersing the TLC plates in 10% H_2_SO_4_ in ethanol, followed by heating to 110 °C.

## Data availability

All phylogenetic data used to produce our results is provided as Supporting Data under the following link: https://data.mendeley.com/preview/xtbwh2k7xc?a=01bb9597-a763-4b51-ab35-e38f36daa29d. All materials of this paper can be provided upon reasonable request. Custom scripts will be made available upon request.

## Acknowledgements

We gratefully acknowledge the C2RT core facilities at the Institut Pasteur, in particular J. Fernandes from UtechS-PBI (supported by France BioImaging; ANR-10–INSB–04; Investments for the Future) and M. Albert and J-Y. Tinevez at the Image Analysis Hub and the Ultrastructural BioImaging facility supported by from the French Government Programmes N° ANR-10-INSB-04-01 and ANR-10-LABX-62-IBEID. We thank P. Campagne from the Bioinformatics and Biostatistics Hub from the IP-C3BI. Molecular graphics were done with ChimeraX, developed at UCSF with support from NIH (R01-GM129325) and NIAID. This work was supported in part by grants from the Agence Nationale de la Recherche (ANR, France), contracts ANR-18-CE11-0017 (P.M.A.) and ANR-21-CE11-0003 (A.M.W.), ANR-24-CE11-4058 (A.M.W), Fondation pour la Recherche Médicale, FRM, EQU202303016284 (P.M.A.), and by institutional grants from the Institut Pasteur, the CNRS, and Université Paris Cité. J.P. was funded through the AMX program from the Ecole Polytechnique. A.S. was part of the Pasteur - Paris University (PPU) International PhD Program, funded by the European Union’s Horizon 2020 research and innovation programme under the Marie Sklodowska-Curie grant agreement No 665807.

## Author contributions

AMW, JP, and PMA designed the research. JP, MM, AS, MBA carried out molecular and cell biology experiments. JP performed cellular imaging and analysis. CT performed SEM experiments. NB and CSA performed and analyzed the TLC experiments. NB, YB and EL provided and synthesized the 2-FIC5Tre probe. DM performed the phylogeny and sequence analyses. JP and AMW wrote the paper. All authors edited the paper.

