## Supplementary Information for "Functional dissection of Wag31 domains for septal recruitment and polar distribution during the cell cycle"

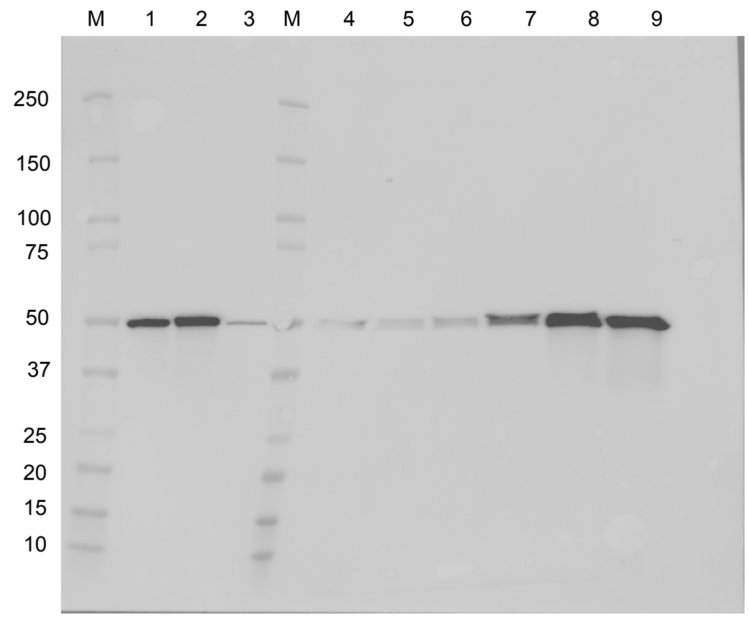


**Supplementary Figure S1**. *Cglu*_P$_{ino}$-*wag31* complementation. Western blot of whole cell extracts from *Cglu* (1), *Cglu* + empty plasmid (2) and *Cglu*_P$_{ino}$-*wag31* (3) after overnight growth in minimal medium. Depleted *Cglu*_P$_{ino}$-*wag31* complemented with P$_{gntk}$-wag31 plasmid in 4% sucrose after 0 (4) and 3 (5) and 6 (6) hours and in the presentce of 1% gluconate (overexpression) after 0 (7) and 3 (8) and 6 (9) hours. Wag31 levels were revealed using anti-Wag31$_{1\text{-}61}$ antibody. Lanes M contain molecular weight markers.

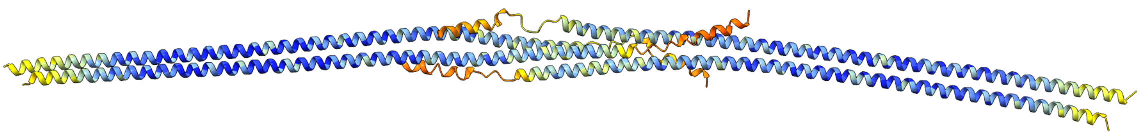

**Supplementary Figure S2**. AF model of the C-terminal tetramerization interface of Wag31.

**Table S1.** Bacterial strains, plasmids and primers used in this study, with descriptions and references.

| **Strains** | **Characteristics** | **Reference** |
|---|---|---|
| ***E. coli*** | | |
| DH5$\alpha$ | F- endA1 Φ80dlacZΔM15 Δ(lacZYA-argF)U169 recA1 relA1 hsdR17(rK–mK+) deoR supE44 thi-1 gyrA96 phoA λ–; strain used for general cloning procedures | [1] |
| CopyCutter EPI400 | F- mcrA Δ(mrr-hsdRMS-mcrBC) Φ80dlacZΔM15 ΔlacX74 recA1 endA1 araD139 Δ(ara, leu)7697 galU galK λ- rpsL (StrR) nupG trfA tonA pcnB4 dhfr; strain used for general cloning procedures | [2] |
| BL21(DE) | F- ompT hsdSB(rB–mB–) gal dcm (DE3); host for protein production | [3] |
| ***C. glutamicum*** | | |
| *Cglu* (ATCC 13032) | Biotin-auxotrophic wild type | [4] |
| *Cglu*_$P_{ino}$-*wag31* (sUMS_214) | *myo*-inositol dependent *wag31* silencing strain. ATCC 13032 with insertion of a terminator and $P_{ino}$ promoter to silence *wag31* (*cg2361)* expression. Repressible in the presence of *myo*-inositol | This work |

| **Plasmids for *C. glutamicum wag31* conditional depletion** | | **Reference** |
|---|---|---|
| *pK19mobsacB* | KanaR; plasmid for allelic exchange in *C. glutamicum*; (pK18 oriVEc, sacB, lacZα) | [5] |
| *pk19-P3323-lcpA* | KanaR; pK19mobsacB derivative. Used as a PCR template to amplify a transcriptional terminator and the promoter of cg3323 ($P_{ino}$) | [6,7] |
| *pK19-*$P_{ino}$*-wag31* | KanaR; pk19mobsacB derivative containing 500 bp upstream-region of *wag31*, a transcriptional terminator, the $P_{ino}$ promoter and 500bp of the *wag31* coding region | This work |

| **Plasmids for recombinant protein expression in *C. glutamicum*.** | | **Reference** |
|---|---|---|
| *pTGR5* | KanaR; *E. coli/C. glutamicum* shuttle vector for regulated gene expression of EGFP under control of tac promoter (Ptac lacI ColE1 oriVEc pGA1 oriVCg) | [8] |
| pUMS_3 | KanaR; pTGR5 derivative in which *Ptac* was exchanged by *PgntK* promoter to control the expression of the EGFP protein | [9] |
| pUMS_17 | KanaR; pUMS3 derivative for expression of Wag31 under control of *PgntK* promoter | This work |
| pUMS_25 | KanaR; pUMS3 derivative for expression of Wag31-mNeonGreen (C-terminal tag) under control of *PgntK* promoter | This work |
| pUMS_21 | KanaR; pUMS3 derivative for expression of mNeonGreen-Wag31 (N-terminal tag) under control of *PgntK* promoter | This work |
| pUMS_175 | KanaR; pUMS3 derivative for expression of $Wag31_{LBD}$ (residues 1-61) fused to mNeonGreen (C-terminal tag) under control of *PgntK* promoter | This work |
| pUMS_176 | KanaR; pUMS3 derivative for expression of $Wag31_{CCD}$ (residues 62-365) fused to mNeonGreen (C-terminal tag) under control of *PgntK* promoter | This work |
| pUMS_254 | KanaR; pUMS3 derivative for expression of $Wag31_{mut}$, carrying the single mutations I18D, K20S, R21S, fused to mNeonGreen (C-terminal tag) under control of *PgntK* promoter | This work |

| | | |
|---|---|---|
| pUMS_255 | KanaR; pUMS3 derivative for expression of Wag31$_{LBD_mut}$, carrying the single mutations I18D, K20S, R21S, fused to mNeonGreen (C-terminal tag) under control of *PgntK* promoter | This work |

| **Oligonucleotide** | **Sequence 5' -->3' and properties[a]** |
|---|---|
| **Plasmids for *C. glutamicum* strain construction** | |
| ***pK19-P$_{ino}$-wag31*** | |
| P7_JP | TGTTGTGTGGAATTGTGCCGCTAGGTAATGTGCGC |
| P8_JP | TTGCGGATTCCCTTCGATTTAACGG |
| P9_JP | GAAGGGAATCCGCAAATAAAACGAAAGGCTCAGTCGAAAGAC |
| P10_JP | TGGAGTCAACGGCATCTAAAATTTCTCCTCTTAAAAAGATAACGGCC |
| P11_JP | ATGCCGTTGACTCCAGCTGATGT |
| P12_JP | AATTGTTATCCGCTCAGTCCACATTTGCAGCACCTGTAGC |

| | |
|---|---|
| **Plasmids for recombinant protein expression in *C. glutamicum*** | |
| ***pUMS_17*** | |
| P178_MM | ATGGTCTTATCCTTTCTTTGGTGGC |
| P179_MM | AGCGGCCGCTTAAGGTAC |
| P216_MM | CCAAAGAAAGGATAAGACCATATGCCGTTGACTCCAGCTGATGT |
| P218_MM | CGGTACCTTAAGCGGCCGCTTTACTCACCAGATGGCTTGTTG |
| ***pUMS_25*** | |
| P178_MM | ATGGTCTTATCCTTTCTTTGGTGGC |
| P179_MM | AGCGGCCGCTTAAGGTAC |
| P214_MM | CTCGAGGGATCTGGCCAGGGACCGGGCTCAGGCCAAGGAAGCGGCATGGTGTCCAAGGGCGAAGAG |
| P215_MM | GAATTCGGTACCTTAAGCGGCCGCTTTACTTGTACAGTTCATCCATGCCCATCACATCGGTGAATG |
| P216_MM | CCAAAGAAAGGATAAGACCATATGCCGTTGACTCCAGCTGATGT |
| P217_MM | CCCTGGCCAGATCCCTCGAGCTCACCAGATGGCTTGTTGTTG |
| ***pUMS_21*** | |
| P178_MM | ATGGTCTTATCCTTTCTTTGGTGGC |
| P179_MM | AGCGGCCGCTTAAGGTAC |
| P190_MM | CAAAGAAAGGATAAGACCATATGGTGTCCAAGGGCGAAG |
| P191_MM | GCCAGATCCCTCGAGCTTGTACAGTTCATCCATGCCC |
| P212_MM | ACAAGCTCGAGGGATCTGGCCAGGGACCGGGCTCAGGCCAAGGAAGCGGCATGCCGTTGACTCCAGCTGATGT |
| P218_MM | CGGTACCTTAAGCGGCCGCTTTACTCACCAGATGGCTTGTTG |

| | |
|---|---|
| ***pUMS_175*** | |
| P13_JP | GGCAACCTGCGCCTCTAGCTCT |
| P14_JP | GCGCAGGTTGCCCTCGAGGGATCTGGCCAGGGAC |
| ***pUMS_176*** | |
| P15_JP | CATATGGTCTTATCCTTTCTTTGGTGGCGT |
| P16_JP | TAAGACCATATGGGTGGTACTTCTTCCGCTGCTAGTT |
| ***pUMS_254 and pUMS_255*** | |
| P39_JP | CCTGACGGCTCCAGTGGCTACAACGAAG |
| P40_JP | CACTGGAGCCGTCAGGCGGCTTATTAAAAGC |

[a]Overlaps for Gibson assembly or site-directed mutations are highlighted in red
